# Acoustic Coupling for Double-Blind Human Low-Intensity Focused Ultrasound Neuromodulation

**DOI:** 10.1101/2025.10.02.680055

**Authors:** Aditya Kapoor, Andrew Strohman, Yunruo Ni, Gabriel Isaac, Jason Raymond, Wynn Legon

## Abstract

**Objective:** Low-intensity focused ultrasound (LIFU) is a non-invasive neuromodulation technique with the potential for precise targeting of small and deep human brain circuitry. To ensure rigorous and reproducible effects, it is necessary to implement double-blinding to mitigate potential experimental biases. This study innovates on previous ultrasound coupling methods by embedding a 3D-printed thermoplastic disc in a high-density gel-polymer matrix to create identical gel-plastic coupling devices that either transmit (verum) or block (sham) ultrasound.

**Methods:** We evaluated varying thicknesses (1.5 mm, 2,0 mm, 2.5 mm) and infill densities (25%, 50%, 75%, and 100%) of Acrylonitrile Butadiene Styrene (ABS) for insertion loss and effects on beam characteristics across relevant human neuromodulation frequencies from 0.2 – 1 MHz.

**Results:** ABS at 1.5 mm thickness and 50 % infill had the lowest insertion loss at 0.50 MHz of 0.9 ± 0.04 dB demonstrating acoustic transparency suitable for a verum device. For the sham device, we introduced an internal air gap that produced 31 dB insertion loss at 0.50 MHz. The verum device demonstrated minimal effects on the beam shape with a radial peak shift of 0.3 ± 0.2 mm and an average axial shift of 0.5 mm.

**Discussion:** This novel gel-plastic coupling device can be tailored to most LIFU transducer shapes and is designed to support double-blind protocols by maintaining visual and tactile indistinguishability while controlling transmitted energy. This method provides a cost-effective and easily implementable solution for double-blinding in human LIFU studies to improve the reliability and rigor of experimental outcomes.

## INTRODUCTION

Low-intensity focused ultrasound (LIFU) is an innovative approach for human non-invasive neuromodulation that uses focused mechanical energy to modulate neural activity with high spatial precision^1,2^. Studies in humans have demonstrated LIFU to influence neural activity across multiple brain regions, including superficial regions like the primary motor^3,4^, visual^5,6^, and somatosensory cortices^2,7,8^ and deeper structures like the insula^9–12^, cingulate cortex^13,14^, thalamus^15^, and putamen^16^. While the precise mechanism underlying LIFU-induced neuromodulation remains under investigation, evidence suggests that ultrasound exerts mechanical forces on cell membranes, activating mechanosensitive ion channels ^1,17^. Unlike other non-invasive brain stimulation techniques such as transcranial electrical stimulation (TES) and transcranial magnetic stimulation (TMS), LIFU enables targeted modulation of both cortical and deep subcortical structures with millimeter-scale resolution and adjustable focal depth^1,20^.LIFU is now being extensively investigated for its therapeutic potential in clinical settings due to its unique capacity to non-invasively target small and deep brain targets implicated in neurological and neuropsychological diseases such as pain^9,11,13,21^, substance use disorders^22,23^ and mental health disorders^1,24^.

Unlike TMS, LIFU must be coupled to the scalp for efficient transmission of ultrasound energy from the transducer through the skin and skull and into the brain. To ensure efficient ultrasound energy delivery, a coupling medium is required between the transducer and scalp because ultrasound does not propagate well through air. At 20 °C and 1 atm, air’s specific acoustic impedance is ∼ 4.1 × 10^2^ kg·m^-2^·s^-1^ (∼ 412 Rayl), which is orders of magnitude lower (∼ 4000x) than that of water (∼ 1.5 × 10^6^ kg·m^-2^·s^-1^ (∼ 1.5 Rayl)) or human skin (1.7 x 10^6^ kg·m^-2^·s^-1^ (∼ 1.7 Rayl)^25^, resulting in a substantial acoustic mismatch that greatly reduces direct energy transfer. Therefore, an acoustic coupling medium with impedance closer to that of water is required to minimize reflection at the skin interface and enable efficient transmission of ultrasound energy from the transducer into the brain.

In addition to effective coupling, it is essential in ultrasound experiments that both verum and sham delivery are indistinguishable to participants and experimenters. Double-blind designs where neither group knows the treatment assignment are a cornerstone of clinical research, minimizing expectancy and observer bias and thereby increasing the rigor and reliability of study findings. This ensures that any observed effects can be attributed to the intervention rather than external influences^26^. However, in human LIFU studies, achieving effective double-blinding remains challenging due to the lack of coupling media that are identical for verum and sham conditions.

Frequencies in the range of 0.2–1 MHz are commonly used in human neuromodulation due to reduced skull-induced attenuation and the ability to maintain precise focal control^1^. In transcranial applications, ultrasound pulses at these carrier frequencies are often delivered at pulse repetition frequencies (PRFs) that can fall within the human audible range^1,27–29^. PRFs of 500 Hz or 1000 Hz are typical in human LIFU protocols, raising the potential for participants to perceive sound. Strategies to mitigate this include auditory masking^9,13,30,31^, active sham conditions^3,32^, or a combination of both^33^. However, these methods often require experimenter intervention, limiting their suitability for double-blind designs. Identical verum and sham coupling devices that keep all procedures equivalent would help eliminate this potential confound and improve experimental rigor.

Here, we build on our prior work with high-density gel polymers as acoustic coupling media ^34^ by incorporating a 3D-printed thermoplastic layer that either transmits or blocks ultrasound energy to create verum or sham LIFU exposures, respectively. Thermoplastics are attractive for LIFU applications due to their tunable acoustic properties, low cost, and ease of fabrication into custom geometries. Different formulations exhibit distinct attenuation characteristics; for example, Acrylonitrile Butadiene Styrene (ABS) has an attenuation coefficient of 1.2 dB/cm at 0.5 MHz^35^. This property can be exploited to fabricate coupling devices that are visually identical yet acoustically distinct—one design highly transparent to ultrasound for verum conditions, and the other strongly attenuating for sham conditions.

This study evaluates Acrylonitrile Butadiene Styrene (ABS) as a potential coupling medium for verum and sham LIFU applications. We examined how variations in 3D printing parameters— specifically thickness and infill density—affect acoustic insertion loss. Material performance was assessed at a fundamental frequency of 0.5 MHz, commonly used in human LIFU experiments^1^, and across a broader range of 0.2–1 MHz relevant to other protocols. We also investigated whether ABS alters ultrasound beam geometry, and tested the feasibility of embedding ABS within a high-density gel polymer^34^ to create a stable interface between the transducer and scalp.

## METHODS

### Experimental Overview

We tested Acrylonitrile Butadiene Styrene (ABS; Ultimaker ABS Black, 2.85 mm filament) samples printed as circular disks at 3 different thicknesses: 1.5 mm, 2 mm, and 2.5 mm. These thicknesses were selected because they fit within 5 mm and 10 mm thick gel polymers that are commonly used in our human LIFU studies^9,11,13,34^. We also evaluated four infill densities (25%, 50%, 75%, and 100%). Infill density refers to the percentage of material inside a 3D-printed structure that affects the integrity of the 3D printed form^36^ and may also affect acoustic insertion loss. The objective was to identify the combination of thickness and infill density of ABS that had the lowest acoustic insertion loss combined with minimal beam aberration. After determining the optimal combination, variability testing was conducted using four independently printed samples to determine stability and reproducibility of the insertion loss. Following verification of consistency between samples, the thermoplastic samples were integrated in a gel-polymer^34^ and further empirically tested for insertion loss and beam deviation. Note: the combination of the ABS with the gel-polymer will be referred to as a ‘gel-plastic device’. Variability testing was performed on the gel-plastic devices to confirm that insertion loss and beam aberration remained consistent across multiple samples. Upon verification of a consistently functioning verum gel-plastic device, a hollow ABS thermoplastic sample was designed by inserting an air gap in otherwise-identical verum ABS sample for testing for the sham condition. Sham thermoplastic samples were characterized both in standalone form and as integrated gel–plastic devices. By inserting an air gap in the original verum ABS sample, the visual and tactile properties of both conditions were maintained, allowing for double-blind experimental designs.

### ABS Thermoplastic Devices

#### Printer and Slicer Settings

We tested ABS based on previous work that demonstrated an acoustic attenuation of 6.6 dB/cm at 1.1 MHz^35^. A 3D printer (Ultimaker-S5, firmware version 8.3.0, hardware type ID 9051-0) was used to make all the prints. Slicer settings were taken from the vendor-supplied ABS material profile in Ultimaker Cura (version 5.10), which loads manufacturer-validated defaults (extrusion and bed temperatures, fan control, layer height, speeds) for this printer–material combination. We used the default settings to ensure stable extrusion and dimensional accuracy. The only parameter intentionally varied for experiments was infill density (25%, 50%, 75%, 100%).

#### Sample Geometry and Mounting

ABS samples were printed as 60 mm-diameter discs to provide a stable handling margin (for handling in the acoustic testing tank (see **Figure 1A & B**) and to cover both the 45 mm active element and the surrounding transducer housing (see *Single Element Focused Transducer)*. The added diameter allowed the sample to be supported above the exit plane without any clips or fixtures intruding into the ultrasound beam path. For hydrophone measurements, the thermoplastic sample was positioned and held perpendicular with the principal direction of wave propagation at a distance of 2.5 mm from the exit plane (see **Figure 1A**). The gel-plastic devices contained a 45 mm diameter ABS thermoplastic disc which matched the transducer’s active diameter and could be completely embedded within the gel-polymer that had a 55 mm diameter (**Figure 1C & D**). This gel-plastic device size provided the optimal dimensions for coupling of this specific transducer with the gel-plastic device (**Figure 1E**).

**Figure 1.**
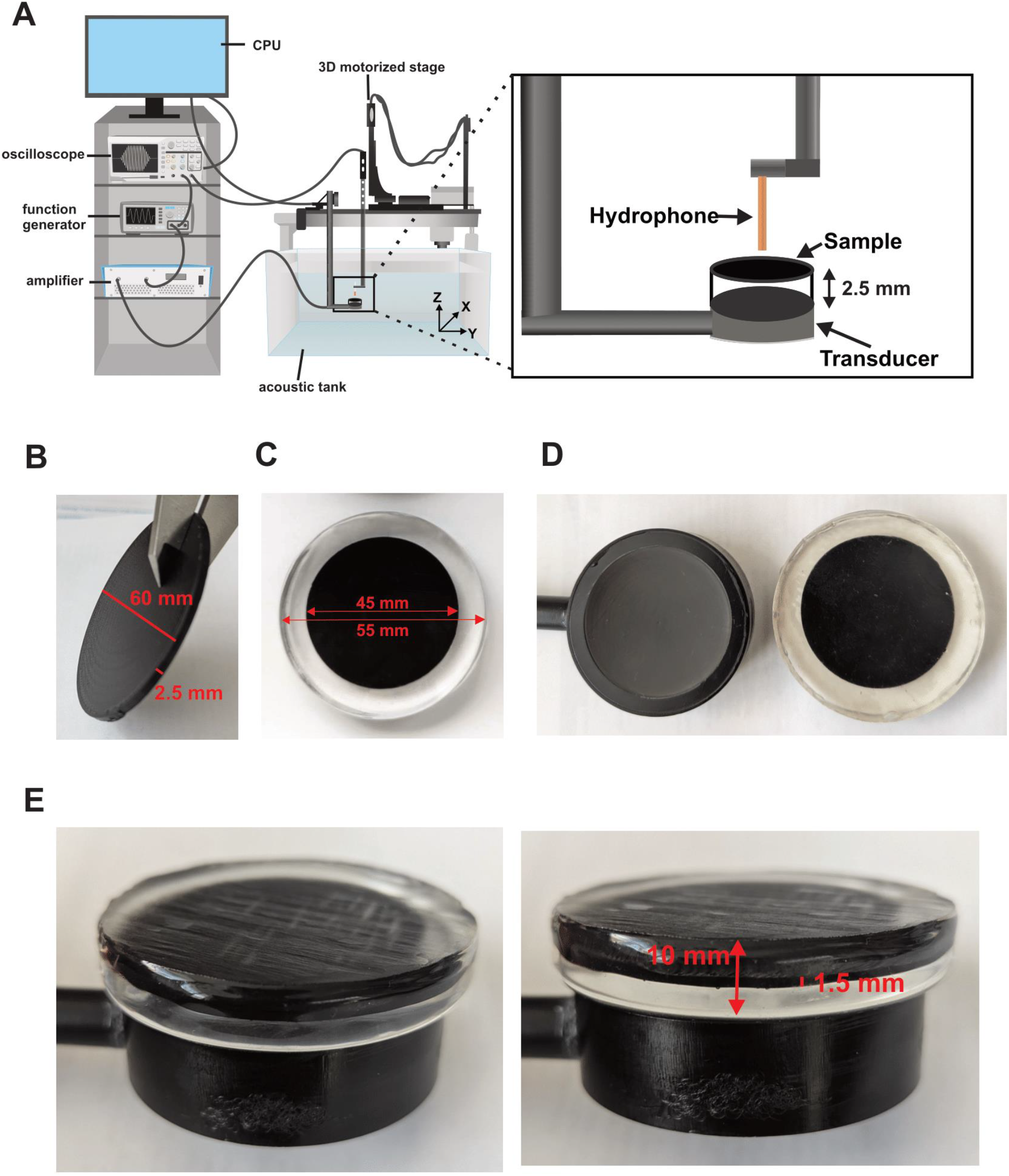
Experimental setup & example ABS discs and gels. **A**. Schematic of the acoustic test tank showing function generator, RF amplifier, oscilloscope, control computer, and a 3D motorized stage. The blow-up demonstrates the positioning of the thermoplastic or gel-plastic device relative to the transducer and hydrophone. **B**. Photograph of 60 mm ABS disc for initial thermoplastic testing. **C**. Photograph of 45 mm ABS disc embedded in 55 mm gel. **D**. Photograph of transducer next to gel-plastic device to show relative sizes. **E**. Photographs of gel-plastic device on transducer used for testing.

#### Parametric Testing of Thermoplastic Samples

A full parametric design approach was used, systematically varying ABS thicknesses (1.5 mm, 2 mm, and 2.5 mm) and infill densities (25%, 50%, 75%, and 100%) resulting in a total of 12 tested samples.

#### Thermoplastic & Gel-Plastic Device Variability Testing

The ABS configuration (thickness and infill density) exhibiting the lowest insertion loss and minimal beam distortion was chosen as the verum condition. The sham condition was subsequently defined by selecting the corresponding ABS air-gap configuration at the same thickness. These sham and verum samples were tested for consistency across four prints. Four devices for each of verum and sham were additionally tested to ensure consistency across prints and device-manufacturing process.

#### Air-Gap (Sham) Thermoplastic Samples

Hollow thermoplastic samples were designed by incorporating an internal shell structure reinforced by five cylindrical pillars. These pillars were positioned along the central axis of the sample, with a diameter of 5 mm and a 9 mm spacing between the center of each pillar. This design ensured that the air gaps were maintained while preserving structural stability during handling and insertion into the gel polymer. The height of the air gap was 0.5 mm for the 1.5 mm thermoplastic sample.

### Integrated Gel-Plastic Device Fabrication

Gel-plastic devices were fabricated using gel curing, heating, and degassing procedures previously described by Strohman et al. (2023)^34^, with modifications made here specifically to accommodate embedding the thermoplastic samples. Briefly, a low-density gel polymer (Penreco “low” density gel wax; Supplier: Amazon, ASIN: B0BK7W1FJL) was heated to its melting temperature of 82.2°C (180°F) in a stainless-steel pot and degassed at 762 mmHg for 10 minutes in a 1.5-gallon vacuum chamber (Orion Motor Tech [OMT], Lake Forest, CA, USA). After degassing, the gel was reheated to 82.2 °C then poured into preheated 10 mm deep, 55 mm diameter silicone molds. The 45 mm thermoplastic sample was positioned centrally between two gel layers, each layer’s thickness adjusted so the final combined thickness of the gel-plastic device was precisely 10 mm (see **Figure 1E**). For example, a 1.5 mm thick thermoplastic disc would be sandwiched between two gel layers, each measuring 4.25 mm. Precision calipers were used to ensure accurate gel layer measurements. Immediately after pouring the first gel layer, the heat gun was set to low heat, and the thermoplastic sample was preheated separately (prior to placement in the gel) for 5 seconds on each side. The preheated sample was then placed at the center of the first gel layer.

After placement of the thermoplastic sample on the first gel layer, a second gel layer was poured on top, fully encapsulating the sample and completing the gel-plastic device formation. To prepare this second layer, the gel was reheated to 93.3 ± 1°C, rested for 2 minutes, and degassed again for 10 minutes. Before pouring the second (top) gel layer, each mold—already containing the first gel layer and thermoplastic sample—was preheated for 20 seconds using low heat to help bonding of the two gel layers. This additional reheating and degassing step is a modification to the previously published process to ensure optimal layer bonding. Devices were left to cure for 45 minutes at room temperature. After curing, any remaining surface bubbles were removed by applying low heat with a heat gun.

### Hydrophone Pressure Measurements and Beam Profile Measurements

All measurements were conducted in an acoustic testing tank (Ultrasonic Measurement System V3 (UMS3), Precision Acoustics Ltd, Dorchester, UK) filled with demineralized, degassed, and filtered water. Waveforms were generated by a function generator (AFG3022C; Tektronix, Beaverton, OR, USA) and amplified using a 100-Watt class A, linear (20 kHz – 12 MHz) amplifier (2100L; Electronics and Innovation, Rochester, NY, USA). A needle hydrophone (HNR-0500, Onda Corp., Sunnyvale, CA, USA), positioned on an automated stage was used to record the acoustic pressure of the transducer. Waveforms were displayed and recorded with an oscilloscope (Keysight Technologies DS0X3012T; El Segundo, CA, USA) and UMS3 (Precision Acoustics Ltd, Dorchester, UK) software. Further analysis was performed using a combination of the post-processing software (Precision Acoustics Ltd; Dorchester, UK) and custom scripts written by WL in MATLAB® (The MathWorks, Inc.; Natick, MA, USA.).

#### Single Element Focused Transducer

A 0.5 MHz single-element spherically focused transducer (H-281, Sonic Concepts Inc., Bothell, WA, USA) was used for testing. The transducer has a 45 mm diameter emitting surface and a radius of curvature of 45 mm (f-number = 1), with a spherical concave cap creating an offset of approximately 6 mm between the exit plane and the geometric base of the transducer, resulting in an effective focal length of 39 mm from the transducer’s exit plane. The transducer concavity was filled with permanent coupling media provided by the manufacturer.

#### Empirical Measurements

##### Line scans

Single-plane line scans were acquired along X, Y, and Z axes using a 25-cycle, 500 kHz pulse at 100 mVpp input voltage. Prior to formal testing, the focal point was located in free-water by iterative X/Y/Z searches with a resolution of 0.1 mm. This co-ordinate was set as [0, 0, 0] mm in X, Y and Z axes and all line scans were made relative to this point. Lateral (X & Y) line scans were acquired over ±4.0 mm about the free-water focal coordinate using the 0.1 mm sampling. The axial line scan consisted of a contiguous 50.0 mm sweep (500 steps at 0.1 mm) ± 25 mm from the free-water focal co-ordinate beginning 1mm off set from the sample. For conditions with the gel-plastic device the standoff height of the hydrophone at its nearest point to the device was 10 mm.

##### Voltage Sweeps

Voltage sweeps were performed using a 25-cycle pulse at 500 kHz with the hydrophone positioned at the peak focal point determined from the free water measurements. Input voltages into the transducer ranged from 20 to 250 millivolts peak-to-peak (mVpp) in 10 mV increments that resulted in peak-to-peak pressure range of 160 kPa to 1930 kPa.

### Analysis

#### Pressure

Insertion loss was calculated relative to the same-day free-water measurement at the focal point (co-ordinate [0, 0, 0]) and reported as the mean ± SD over the pressure range tested (160 kPa – 1930 kPa). Insertion loss was calculated in decibels (dB) by comparing the mean acoustic pressure between free water and sample/device conditions, defined as:

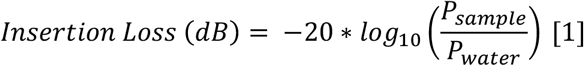

Percent pressure transmission was also calculated from insertion loss as follows:

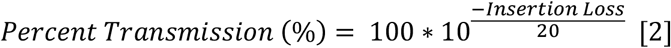

Percent pressure attenuation was calculated as follows:

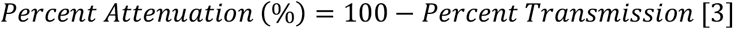

#### Line scans

To determine any shift in each of the X, Y and Z axes, we calculated the change in the position of peak pressure between the free-water scan and the scan through the ABS disc and the gel-plastic device. We also calculated the radial displacement using the following formula:

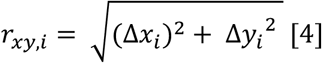

Spatial beam characteristics were quantified by measuring two width metrics along each axis: the full width at half maximum of acoustic pressure (FWHM acoustic pressure, –6 dB width) and the full width at –3 dB acoustic intensity (–3 dB width, corresponding to ∼71 % of peak pressure. Each metric was calculated from the line scans in the X, Y, and Z axes to assess translation of the focal point or beam expansion or contraction introduced by the samples or devices. Of note, the HNR-0500 needle hydrophone has a 0.5 mm active element and thus measurements at 0.1 mm represent a spatial average over this diameter.

### Broadband Insertion Loss

Broadband insertion loss measurements were conducted by placing the sample between two transducers in a tank (60 cm × 15 cm × 20 cm) filled with demineralized, degassed, and filtered water. A transmit transducer (ULTRAN GS500-D25-P50-MR, 25 mm diameter, 50 mm focal length, 500 kHz center frequency, S/N 340373) and a receive transducer (AEROTECH Gamma, 0.75’’ diameter, 10-MHz center frequency, S/N A29617) were co-aligned and positioned approximately 340 mm apart. An ultrasound pulser-receiver (DPR300, JSR Ultrasonics, Pittsford, NY) was used to generate the electrical excitation pulse and amplify the received ultrasound signals, which consisted of 2 acoustic cycles centered at 500 kHz. Signals were captured using a digitizing oscilloscope (Keysight DSOX1204G; 8 bits, 1 GSPS sampling rate) and the frequency-dependent insertion loss of the sample was determined by comparing the Fourier-transformed frequency spectrum of voltage-time waveforms acquired with and without the sample. Results are reported at 20-kHz intervals in the frequency range 0.2 MHz to 1.0 MHz corresponding to the −20 dB bandwidth of the transmit-receive system.

## RESULTS

### Free Water Empirical Measurements

Routine daily calibrations in free-water confirmed that the transducer produced a highly linear pressure–drive relationship (slope = 7.64 ± 0.06 kPa mVpp^−1^, R^2^ ≥ 0.999). Across four calibration sessions collected on different days the coefficient of variation of peak pressure was ≤ 1.6 %. Pressures were 164 ± 1 kPa at 20 mVpp, 795 ± 12 kPa at 100 mVpp, and 1924 ± 31 kPa at 250 mVpp. See **Figure 2A**.

**Figure 2.**
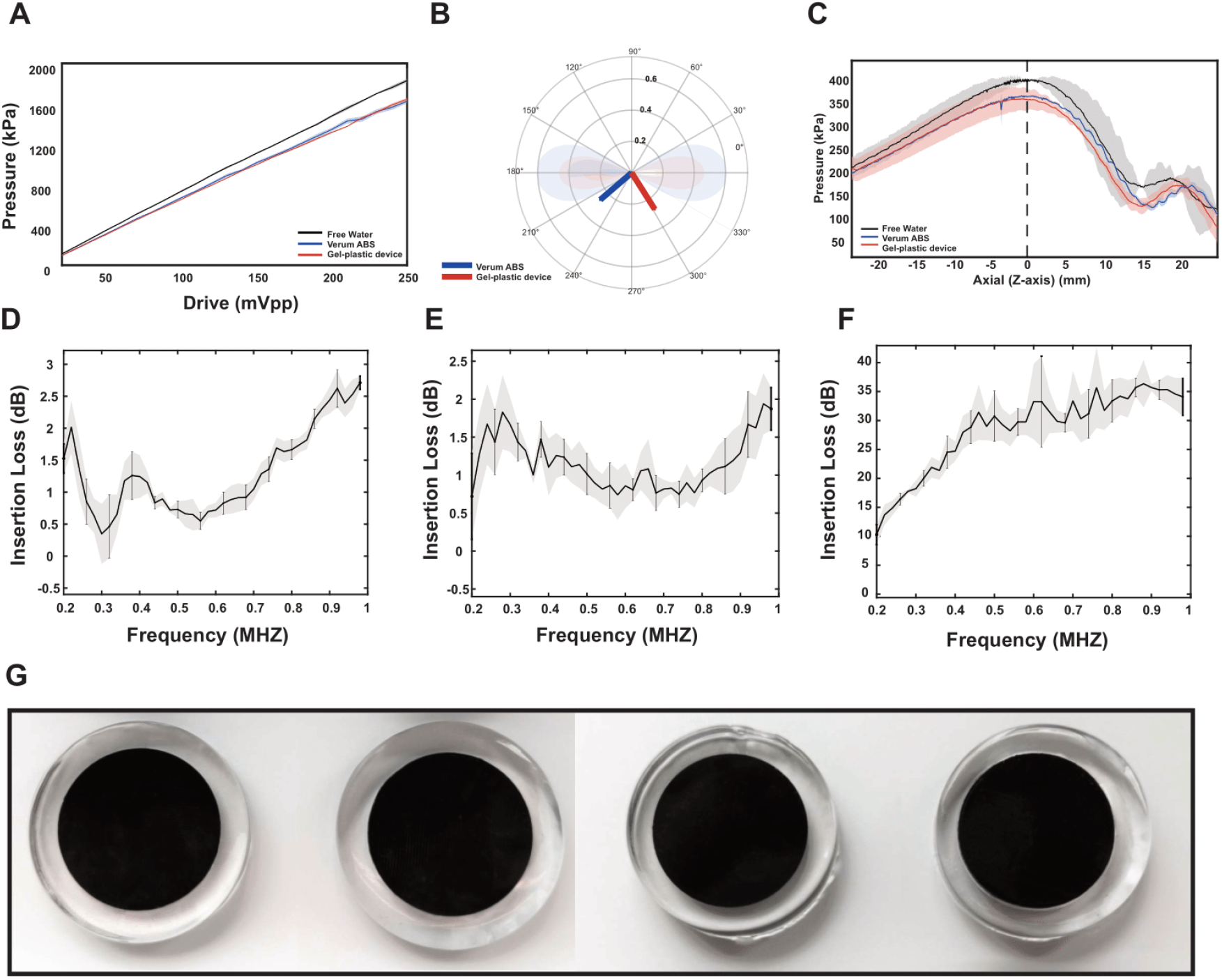
Line scan and pressure insertion loss. **A**. Pressure–drive function in free water (black), the verum ABS disk (blue; 1.5 mm, 50% infill density), and the gel-plastic device (red). Data are shown as group means with ±SD error bars. **B**. Polar plot of lateral peak shift vectors for verum ABS (blue) and gel-plastic devices (red), with free water at the origin. Arrows indicate mean vector direction and length; shaded wedges and rings indicate ±1 SD in angle and radial displacement, respectively. **C**. Axial line plots (Z-axis) in free water (black), the verum ABS disk (blue), and the gel-plastic device (red), plotted as group means ± SD. Vertical dashed line (0) represents peak pressure for the verum ABS conditions that was set as zero (0) on the x-axis. The transducer would be located on the right axis roughly 10 mm to the right for a focal length from the exit plane of ∼ 40 mm. **D**. Mean ± SD broadband insertion loss through the 1.5 mm, 50% infill density sample. **E**. Mean ± SD broadband insertion loss through the gel-plastic device.**F**. Mean ± SD insertion loss through the sham gel-plastic device. **G**. Top-view photographs of sham and verum devices side-by-side for comparison.

Lateral beam profiles were highly consistent across days. Mean ± SD FWHM (−6 dB pressure) for X, Y and Z axes was: 5.0 ± 0.1 mm, 5.1 ± 0.1 mm and 40.0 ± 0.7 mm. The mean ± SD of the −3dB pressure for X, Y and Z was: 3.6 ± 0.1 mm, 3.7 ± 0.1 mm and 25.9 ± 0.1 mm. The focal point as assessed by radial displacement was 0.2 ± 0.1 mm. See **Figure 2 B & C**.

### Thermoplastic Thickness and Infill Density Testing

Insertion loss at 500 kHz was measured for ABS thermoplastic samples across thicknesses of 1.5, 2.0, and 2.5 mm and infill densities of 25, 50, 75, and 100 %. Insertion loss across all tested samples ranged from 0.7–2.0 dB (7.9%-20.4%) with the 1.5 mm 50% infill density demonstrating the lowest insertion loss of 0.7 dB and 2.0 mm thickness and 50% infill density with the highest (2.0 dB); see **Table 1** and **Figure 2A-C**. As such, the 1.5 mm, 50 % infill configuration was selected as the verum thermoplastic sample for subsequent integrated-device testing due to its minimal insertion loss and minimal thickness, allowing for incorporation in various coupling gel thicknesses.

**Table 1.** Insertion loss (dB) at 500 kHz for ABS thermoplastic samples.

| Thickness (mm) | Infill Density (%) | Mean Insertion Loss (dB) |
| --- | --- | --- |
| 1.5 | 25 | 1.8 |
|  | 50 | 0.7 |
|  | 75 | 1.6 |
|  | 100 | 1 |
| 2 | 25 | 1.3 |
|  | 50 | 2 |
|  | 75 | 1.5 |
|  | 100 | 1.6 |
| 2.5 | 25 | 0.9 |
|  | 50 | 0.6 |
|  | 75 | 0.8 |
|  | 100 | 1.2 |

### Verum ABS Testing

#### Pressure

We did subsequent testing of the 1.5 mm, 50% infill density ABS disc. Across 4 independent measurements the mean ± SD insertion loss was 0.8 dB ± 0.1 dB (independent sample insertion loss: 0.9, 0.7, 0.7, 0.9). See **Figure 2A**.

#### Line scans

The FWHM through the 1.5 mm, 50% infill density for X, Y and Z axes was: 5.0 ± 0.1 mm, 5.3 ± 0.1 mm, and 39.0 ± 0.0 mm respectively resulting in respective Δ = 0.0, 0.2, 0.0 mm relative to free water. The −3 dB width for X, Y and Z axes was: 4.0 ± 0.1 mm, 3.9 ± 0.1 mm, and 24.0 ± 0.0 mm respectively resulting in respective with Δ = 0.0, 0.2, 0.0 mm relative to free water. The peak-pressure focal point differed by −0.1 ± 0.4 mm in the X-axis and −0.1 ± 0.1 mm in the Y-axis, resulting in a mean ± SD radial displacement of 0.2 ± 0.1 mm. The location of the axial Z peak pressure was 38.5 ± 0.1 mm, corresponding to ΔZ = −0.45 mm relative to same-day free water (39 mm). The mean ± radial shift of the focal point was 0.4 ± 0.1 mm. See **Table 2 & Figure 2 B-C**.

**Table 2.** Verum ABS thermoplastic (1.5 mm, 50% infill) characteristics.

| Parameter | Axis | Free Water (mm) | Verum mean ± SD (mm) | $\Delta$ (verum – free water) (mm) |
| --- | --- | --- | --- | --- |
| -6dB | X | 5 | 5.0 ± 0.1 | 0 |
| | Y | 5.1 | 5.3 ± 0.1 | 0.2 |
| | Z | 37 | 36 ± 0.0 | 0 |
| -3dB | X | 4 | 4.0 ± 0.1 | 0 |
| | Y | 3.7 | 3.9 ± 0.1 | 0.2 |
| | Z | 24 | 24 ± 0.0 | 0 |
| Focal point | X | 0 | -0.1 ± 0.4 | -0.1 |
| | Y | 0 | -0.1 ± 0.1 | -0.1 |
| | Z | 39 | 38.5 ± 0.1 | -0.45 |
| Radial Shift | | 0 | 0.4 ± 0.1 | 0.2 ± 0.1 |

#### Broadband Insertion Loss

Insertion loss decreased from 1.5 dB at 0.20 MHz to a minimum of 0.3 dB at 0.30 MHz, then increased to 2.7 dB at 1.0 MHz. At 0.50 MHz, insertion loss was 0.7 ± 0.1 dB (mean ± SD). See **Figure 2D**.

### Effect of Verum gel-plastic device on signal transmission

#### Pressure

At 500 kHz, insertion loss measured across four independently manufactured gel-plastic devices, the mean ± SD was 0.90 ± 0.04 dB (1.0 dB, 0.9 dB, 0.9 dB, 0.9 dB). See **Figure 2A**.

#### Line scans

Line-scan measurements (X, Y, Z) were acquired at 500 kHz for the verum integrated gel-plastic device using the same hydrophone coordinates determined for the same-day free-water focal point. The FWHM (−6dB) for X, Y and Z axes was: 5.1 ± 0.1 mm, 5.1 ± 0.1 mm, and 36 ± 0.4 mm respectively resulting in respective Δ = 0.0, 0.0, 0.0 mm. The −3 dB width for X, Y and Z axes was: 3.5 ± 0.1 mm, 3.6 ± 0.1 mm, and 23 ± 0.3 mm respectively, resulting in respective Δ = −0.1, −0.1, 0.0 mm relative to free water. The mean ± SD peak − pressure location for the X-axis was 0.3 ± 0.2 (X) and −0.1 ± 0.2 mm for the Y-axis. The mean ± SD axial peak was −0.5 ± 0.3 mm from the exit- plane origin. See **Table 3 & Figures 2 B & C**.

**Table 3.** Gel-plastic device characteristics.

| Parameter | Axis | Free Water (mm) | Verum mean ± SD (mm) | $\Delta$ (verum – free water) (mm) |
| --- | --- | --- | --- | --- |
| -6dB | X | 5.1 | 5.1 ± 0.1 | 0 |
| | Y | 5.1 | 5.1 ± 0.1 | 0 |
| | Z | 36 | 36 ± 0.4 | 0 |
| -3 dB | X | 3.6 | 3.6 ± 0.1 | 0 |
| | Y | 3.7 | 3.6 ± 0.1 | 0.1 |
| | Z | 23 | 23 ± 0.3 | 0 |
| Focal point | X | 0 | 0.3 ± 0.2 | -0.3 |
| | Y | 0 | -0.1 ± 0.2 | -0.1 |
| | Z | 39 | 38.5 ± 0.3 | -0.5 |
| Radial Shift | XY | 0 | 0.3 ± 0.2 | 0.3 ± 0.2 |

Using the lateral radial metric, the mean ± SD focus was displaced by 0.3 ± 0.2 mm relative to same-day free water (see **Table 3**).

#### Broadband Insertion Loss

Broadband insertion loss measurements from 0.20-1.0 MHz showed frequency-dependent insertion loss. The mean insertion loss fell from 1.6 ± 0.3 dB at 0.20 MHz to a minimum of 0.8 ± 0.1 dB at 0.60 MHz, then rose steadily to 2.0 ± 0.2 dB at 0.95 MHz and 2.2 ± 0.2 dB at 1.00 MHz (mean ± SD, n = 4). At 0.50 MHz, insertion loss was 1.0 ± 0.1 dB (mean ± SD). See **Figure 2E**.

### Effect of Sham gel-plastic device on Signal Transmission

#### Pressure

At 500 kHz, insertion loss measured from voltage sweeps (20–250 mVpp) across four independently manufactured integrated gel-plastic devices was 31.8 ± 1.3 dB (mean ± SD; n = 4; individual device values are 33.4, 32.1, 30.2, and 31.4 dB).

#### Line scans

Evaluation of beam profile shifts were not analyzed.

#### Broadband Insertion Loss

Broadband insertion loss measurements from 0.20-1.0 MHz showed average frequency-dependent insertion loss for the 1.5 mm sham gel-plastic devices (n=4). The mean insertion loss increased from 10.3 ± 1.7 dB at 0.20 MHz to 30.8 ± 4.3 dB at 0.50 MHz, rose further to 33.3 ± 6.5 dB at 0.60 MHz, peaked at 35.4 ± 2.5 dB at 0.95 MHz, and then declined slightly to 34.1 ± 3.3 dB at 1.00 MHz (mean ± SD; n = 4). See **Figure 2F**.

See **Figure 2G** for comparison of verum and sham devices in no particular order.

## DISCUSSION

This study evaluated 3D-printed Acrylonitrile Butadiene Styrene (ABS) as a coupling material— alone and integrated within a gel polymer—for suitability as verum and sham coupling media for use in human low-intensity focused ultrasound (LIFU) neuromodulation studies. We assessed how different ABS configurations (thickness and infill density) affects pressure insertion loss and the resultant focused ultrasound beam geometry. We found that the 1.5 mm thickness combined with 50 % infill density had the lowest insertion loss of 0.9 dB at 500 kHz (n=4) with nominal effect on beam characteristics. As such, we further tested this configuration embedded in a gel polymer and found this did not appreciably affect results making this device suitable for a verum coupling medium. We then tested for any variability in the manufacturing process and found a low variability in insertion loss and beam geometry demonstrating that these devices can be made consistently. We did not achieve no loss with our verum device and methods to improve this are ongoing. However, the ∼ 1dB loss is acceptable and can be accounted for. Under linear acoustic propagation—when the transmitting media and device configuration/geometry remain unchanged—attenuation is a fixed proportion of the wave amplitude and accordingly, the pressure ratio (and insertion loss) is independent of drive level over the tested range (160–1930 kPa). For example, with an incident pressure of 200 kPa, the verum device allows through ∼ 89% or ∼ 178 kPa. To achieve the desired 200 kPa after passing through the verum medium, the experimenter could simply use the inverse of this transmission (1/0.89) and multiply it by 200 kPa that results in an initial pressure of 224 kPa to achieve 200 kPa after passing through the verum device. This pressure would then also be used for the sham condition to preserve blinding.

The verum device resulted in minimal beam deviation of ∼ 0.5 mm. This shift is deemed acceptable for human studies as this is considerably smaller than human cortical gyri that have widths of ∼ 10 mm (precentral ∼10 ± 5 mm; postcentral ∼7 ± 3 mm)^37^. This shift is also acceptable for deep brain targets as for example, the ventral intermediate (VIM) nucleus, a common target for high intensity focused ultrasound, measures roughly 4 × 4 × 6 mm (mediolateral ≈ 4 mm)^38^.

While the majority of human neuromodulation studies employ 500 kHz fundamental frequency; frequencies in the range of 0.2 to 1 MHz have been used. As such, we tested insertion loss across this frequency range and found frequency-dependent insertion loss that in general, increased with higher frequencies though this was not linear and demonstrated some variability. Nevertheless, the maximum insertion loss was ∼ 2.5 dB at 1 MPa that could be compensated for as illustrated previously. Due to the non-linear nature of the frequency response, perhaps different thicknesses and infill densities are more efficacious for different frequencies. We did not test this but is a focus of future work. Regardless, we tested the insertion loss at 500 kHz using two different methods and found good agreement demonstrating stable and reliable acoustic properties.

For the sham device we preserved external design parameters (to maintain appearance)— material (ABS), thickness (1.5 mm), infill pattern (50 %), diameter (45 mm), and color—and added an internal air gap stabilized by cylindrical pillars so the sample could withstand heating/curing and feels and looks identical when embedded in the gel. Because the objective of the sham device is to block ultrasound, detailed evaluation of beam-profile shifts was not performed – as under such conditions, lateral/axial peak definitions become dominated by noise and do not reflect a usable focal field. The air gap produced the expected high attenuation (≈31 dB), while visual/tactile indistinguishability supports double blind use. We informally tested visual and tactile similarity of verum and sham devices with individuals in our lab familiar with human LIFU procedures. They were unable to distinguish sham from verum above chance. We have provided photographs of four random (verum, sham) devices for readers to assess visual likeness (Fig 2G).

While the verum device resulted in ∼ 1 dB insertion loss, our design achieved ∼ 30 dB difference between sham and verum conditions which is suitable for double-blind experiments. The integrated verum device provides a cost-effective and easily implemented solution for double-blinding in human LIFU studies. To our knowledge, there is no commercially available alternative that offers this level of experimental control, and the ability to mitigate participant awareness and experimenter bias outweighs the partial loss in transmission efficiency.

### Limitations & Future Work

A potential limitation involves the structural design of the sham devices. The internal pillars were designed with a 5 mm diameter to ensure mechanical stability during the high-heat gel embedding process. However, this pillar size exceeds the wavelength of ultrasound at frequencies above 300 kHz (wavelength in water at 500 kHz is ∼3 mm), introducing acoustic impedance mismatches that can scatter rather than effectively reflect or attenuate the ultrasound wave. Structures larger than the wavelength can more effectively couple the energy across the layer thus diminishing overall insertion loss. In contrast, sub-wavelength features (<1 wavelength) can more effectively disrupt wave propagation through impedance mismatch without introducing unwanted scattering artifacts.

We did not conduct any durability or longevity testing. However, anecdotally, the integrated gel– plastic devices are reusable with careful handling but the gel-polymer will eventually degrade and chip with repeated use due to mechanical stress. The gel-polymer will not however, degrade due to warming from contact with the skin as is common with more viscous coupling mediums such as ultrasound gels. To preserve performance and blinding, we recommend: (i) storing devices in a sealed container when not in use to limit gel dehydration; (ii) avoiding excessive heat exposure; (iii) pre-session visual inspection for microbubbles, delamination, or surface damage; and (iv) a regularly scheduled quick insertion-loss check against the device’s baseline, flagging any drift ≥1 dB for retirement. Devices with visible defects or measurable acoustic drift should be removed from service.

Future work will investigate different ABS thicknesses and infill density combinations for other common frequencies and work to reduce insertion loss to zero. For the sham devices, work will investigate the effect of reducing pillar diameters and arranging them in a regular pattern throughout the sample. This could enhance insertion loss at higher frequencies by promoting more efficient wave disruption while still maintaining mechanical support. However, smaller pillars may compromise the device’s structural integrity during fabrication and handling. Thus, future efforts should focus on optimizing this tradeoff between acoustic performance and mechanical stability to further improve hollow device efficacy.

## Conclusion

This study quantifies the insertion loss and beam aberration of ABS, both alone and when integrated with gel polymers, to evaluate its suitability as a coupling device for rigorous double blind human LIFU neuromodulation. The device offers a cost effective, reproducible way to improve blinding and experimental rigor in human LIFU research and clinical trials.

## ACKNOWLEDGEMENTS

This work was supported by grants from the NIH R21AT012247 & UG3DA059407 and Focused Ultrasound Foundation to WL and from the NIH F30AT013174 to AS. We would also like to thank Jessica Florig & Kathryn Painchaud for assistance in the device manufacturing process.

## DATA AVAILABILITY STATEMENT

Data is available upon request.

